# eMOSAIC: Multi-modal Out-of-distribution Uncertainty Quantification Streamlines Large-scale Polypharmacology

**DOI:** 10.1101/2024.01.05.574359

**Authors:** Amitesh Badkul, Li Xie, Shuo Zhang, Lei Xie

## Abstract

Polypharmacology has emerged as a new paradigm to discover novel therapeutics for unmet medical needs. Accurate, reliable and scalable predictions of protein-ligand binding affinity across multiple proteins are essential for polypharmacology. Machine learning is a promising tool for multi-target binding affinity predictions, often formulated as a multi-modal regression problem. Despite considerable efforts, three challenges remain: out-of-distribution (OOD) generalizations for compounds with new chemical scaffolds, uncertainty quantification of OOD predictions, and scalability to billions of compounds, which structure-based methods fail to achieve. To address aforementioned challenges, we propose a new model-agnostic anomaly detection-based uncertainty quantification method, **e**mbedding **M**ahalanobis **O**utlier **S**coring and **A**nomaly **I**dentification via **C**lustering (eMOSAIC). eMOSAIC uniquely quantifies distribution similarities or differences between the multi-modal representation of known cases and that of a new unseen one. We apply eMOSAIC to a multi-modal deep neural network model for multi-target ligand binding affinity predictions, leveraging a pre-trained strucrture-informed large protein language model. We extensively validate eMOSAIC in OOD settings, showing that it significantly outperforms state-of-the-art sequence-based deep learning and structure-based protein-ligand docking (PLD) methods by a large margin as well as existing uncertainty quantification methods. This finding highlights eMOSAIC’s potential for real-world polypharmacology and other applications.

## 1 Introduction

Drug discovery is highly complex process, taking up to 15 years and costing billions of dollars [1], but its failure rate is extremely high due to lack of clinical efficacy or safety [2]. Many complex diseases such as neurological and mental disorders are multigenic, multi-factorial diseases. Thus, an effective therapy needs to modulate multiple genes [3, 4]. Polypharmacology that uses a single chemical to selectively modulate multiple drug targets has emerged as a new paradigm in drug discovery [5–10]. Screening a library of billions of compounds against multiple drug targets to identify lead compounds and subsequently optimizing their binding affinity selectivity profiles via medicinal chemistry for drug candidates are critical steps in polypharmacology [11]. A giga-scale compound screening is needed for several reasons. First, it can proportionally increase the likelihood of identifying more potent or selective ligands [12–15]. Secondly, the accessibility of analogues of lead compounds in the same chemical spaces can streamline the generation of meaningful structure–activity relationship (SAR) for lead optimization. Finally, large-scale libraries can expand chemical diversity, chemical novelty and patentability of the drug leads [16]. Although DNA-encoded libraries are an effective approach to generate and screen billions of compounds for a single target[17], it has limited chemistries due to DNA conjugation and a high rate of false positives by nonspecific binding of DNA labels. It is envisioned that computational approaches will facilitate large-scale compound screening for polypharmacology[11].

Protein-ligand docking (PLD) is commonly used for polypharmacology screening when the 3D structure of drug target is experimentally determined or computationally predicted [18]. However, PLD suffers from a high rate of false positives due to poor modeling of protein dynamics, solvation effects, crystallized water, and other challenges [19]. The reliability of PLD significantly deteriorates when using predicted structures [20, 21], and is not reliable for virtual screening [22]. Despite the success of AlphaFold2 [23], it can only reliably model half of understudied human proteins whose small molecule ligands are unknown [24]. Moreover, PLD is computationally intensive, taking several seconds to score a protein-chemical pair. Thus, it is not feasible for PLD to screen billions of compounds against multiple targets for polypharmacology.

Recent advances in deep learning (DL) encourage increasing interest in applying DL to drug discovery [25, 26]. Sequence-based DL, an alternative to traditional docking methods, offers significant advantages. Sequence-based drug-target interaction (DTI) prediction, which uses 3D structure information only implicitly, enables fast DTI predictions when the inputs consist solely of a molecular description of the drug represented by a SMILES string or a 2D graph and the amino acid sequence of the target protein [27, 28]. By leveraging protein sequences instead of full 3D structures, we can reduce the computational burden and allow for the rapid evaluation of vast libraries of compounds, making it a more practical and scalable strategy for polypharmacology. However, the generalization power of deep learning methods for protein-ligand interaction predictions remains poor. The chemical space of small organic molecules is astronomical vast. Although the number of possible small organic molecules is approximate 10^60^ [29], only around 10^6^ compounds have annotated protein targets [30–32]. The limited coverage of chemical genomics space make it challenging to train generalizable deep learning models for binding affinity predictions in an out-of-distribution (OOD) scenario for unexplored chemical space [33, 34], in which unseen testing chemicals are significantly different from training data.

Since drug discovery is a high stake process, making decisions based on incorrect predictions can lead to time and resource wastage. Knowing the confidence level of a prediction is crucial, as it allows researchers to make informed decisions about whether to consider or disregard specific drug leads. This necessitates an estimation of a reliability measure for individual predictions. The uncertainty of prediction comes from either the lack of labeled data or data noisiness. Gaussian Process (GP) is one of popular approach to the uncertainty quantification. Several works [35, 36] propose a combined GP and multi-layer perceptron (MLP) approach for various biological tasks. However, the proposed GP+MLP algorithm is computationally intensive and requires the modification of the architecture of predictive models. Zeng and Gilford [37] implement an ensemble of NNs to obtain the uncertainty associated with the predictions for peptide-MHC binding. However, the ensemble-based technique is not as accurate as the GP algorithm for quantifying uncertainty. Along with these methods, Conformal Prediction aims to provide distribution-free uncertainty quantification under data exchangeability assumptions, a weaker version of independent and identically distribution (IID) [38, 39]. Many works have attempted to utilize conformal prediction for various aspects in drug discovery ranging from Quantitative Structure-Activity Relationships (QSAR), Quantitative Structure-Property Relationships (QSPR), pIC50, and toxicity prediction [40–42]. While conformal prediction in the case of classification provides accurate uncertainty quantification, these methods often struggle with efficiency in regression tasks by estimating large confidence intervals. Moreover, the exchangeablility assumption underlying the conformer prediction does not hold in OOD cases.

To overcome the aforementioned challenges in large-scale polypharmacology compound screening, we propose a new model-agnostic anomaly detection-based uncertainty quantification method, **e**mbedding **M**ahalanobis **O**utlier **S**coring and **A**nomaly **I**dentification via **C**lustering (eMOSAIC). eMOSAIC explicitly models multi-modal distributions of embedding space and quantifies the similarity between embeddings of known cases and that of a new one. We apply eMOSAIC to a multi-modal deep learning network model for multi-target ligand binding affinity predictions, leveraging a pre-trained structure-informed large protein language model[43]. Under rigorous OOD benchmark studies, eMOSAIC significantly outperforms state-of-the-art deep learning models for binding affinity prediction. Interestingly, eMOSAIC also demonstrates superiority over PLD in terms of both accuracy and scalability. Furthermore, eMOSAIC outperforms existing uncertainty quantification methods, demonstrating its potential for other machine learning applications. Thus, eMOSAIC represents a significant advance in deep learning applications to drug discovery.

In summary, our contributions encompass both methodology development and translational applications:

1. We introduce eMOSAIC, a novel OOD multi-modal uncertainty quantification method that outperforms state-of-the-art methods and may have broad applications in addressing the OOD challenge.
2. By leveraging a pre-trained structure-informed protein language model, we demonstrate that eMOSAIC outperforms state-of-the-art deep learning methods in protein-ligand binding affinity prediction and polypharmacology compound screening.
3. Through rigorous benchmark studies, we show that sequence-based eMOSAIC significantly surpasses the existing structure-based compound screening methods dominant in drug discovery in terms of accuracy, reliability, and scalability.

## 2 Results

### 2.1 Overview of eMOSAIC

Figure 1 provides an overview of eMOSAIC. Similar to state-of-the-art uncertainty quantification methods such as conformal prediction, eMOSAIC utilizes the training data to train a secondary model for uncertainty estimation alongside the task-specific base models. However, unlike conventional methods, eMOSAIC generalizes well to OOD data. The training of eMOSAIC involves three main steps. First, the embeddings of training examples from a trained task-specific deep learning model are clustered. Second, for a given validation case, the Mahalanobis distances between its embedding and each cluster of training embeddings are calculated, and the absolute residual of its prediction is obtained using the trained model. Finally, eMOSAIC is trained using these Mahalanobis distances as features and the absolute residuals as labels.

**Fig. 1.**
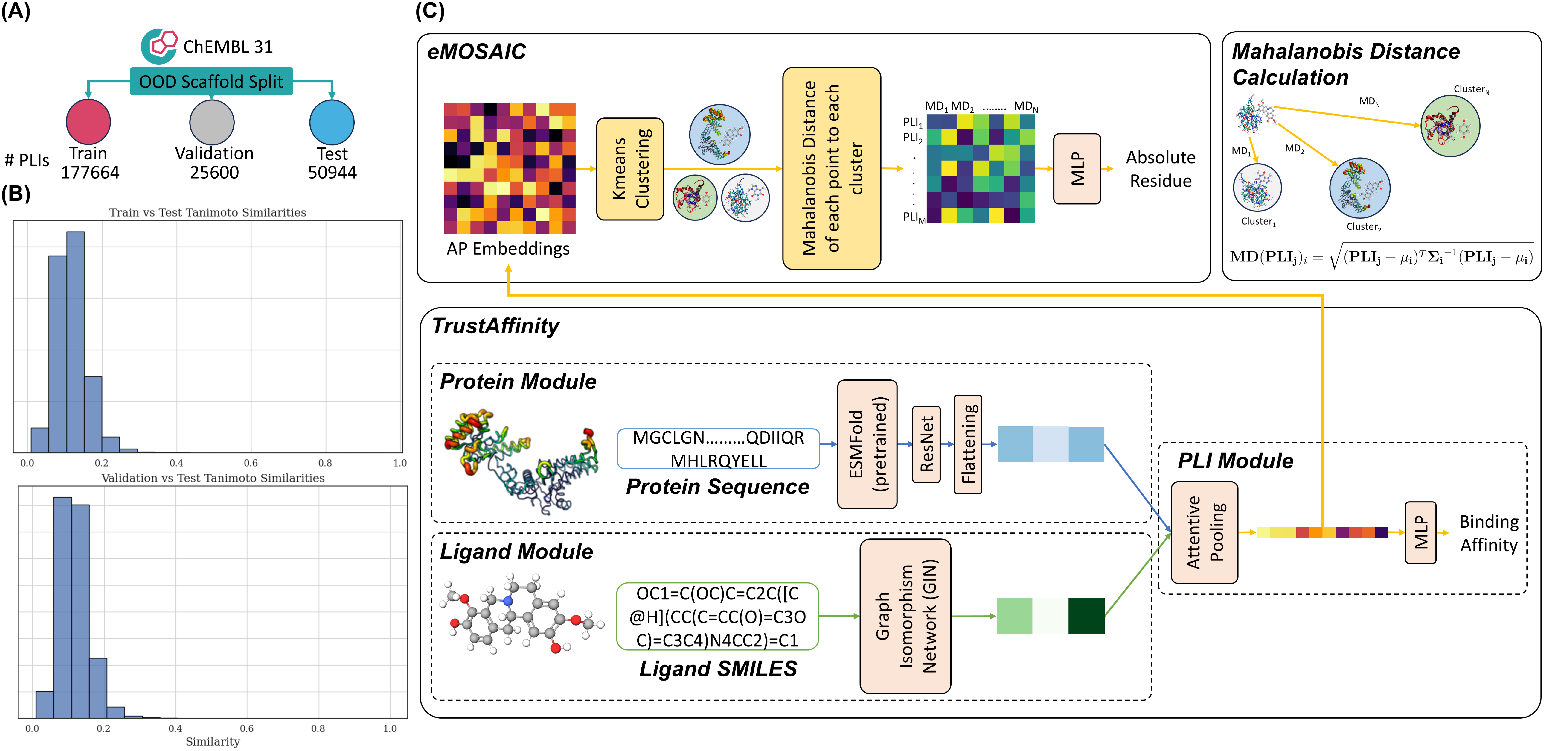
Training data and architecture of eMOSAIC. (A) ChEMBL 31 dataset OOD Scaffold Split to ensure no overlap of scaffolds among the train, test, and validation set. (B) Tanimoto similarity histogram of the test vs validation and test vs train set, respectively. (C) Overall architecture of the pipeline with the binding affinity prediction module, TrustAffinity, and the model agnostic uncertainty quantification module, eMOSAIC.

eMOSAIC is model-agnostic and can be applied to any trained task-specific DL models with embeddings. In this paper, we apply it to a protein-ligand binding affinity prediction model, called as TrustAffinity. The affinity prediction model takes a protein sequence and a ligand SMILES as inputs. Leveraging a pre-trained protein language model ESMFold [43] and a graph isomorphism network (GIN) [44], the embeddings of protein sequences and chemical structures are combined via attention pooling. The generated vectors from the attention pooling are used to predict binding affinities and train eMOSAIC for uncertainty quantification. For simplicity, whenever eMOSAIC is mentioned, it always refers to its application to the TrustAffinity model.

We evaluated the performance of eMOSAIC in a scaffold-based OOD setting, where chemicals in the testing set have different chemical scaffolds from those in the training/validation set mimicking real-life scenario, where we are likely to encounter new potential chemicals which would have not been used for training process of the DL model. For the purpose of comparison, we also evaluate eMOSAIC in the in-distribution setting of random split.

We compared eMOSAIC with state-of-the-art methods in three tasks, 1) Binding Affinity Prediction, 2) Polypharmacology Compound Screening, and 3) Uncertainty Quantification. For binding affinity prediction and polypharmacology screening tasks, we compare eMOSAIC with both sequence-based deep learning models including BACPI [45], DeepDTA [46], DeepPurpose [47] as well as a typical PLD method, AutoDock Vina [48], and deep learning-based docking model KarmaDock [49]. Along with this, we also compare the performance of eMOSAIC with other uncertainty quantification methods including GP-based RIO framework [36], Monte Carlo Dropout [50], and Conformal Prediction [38].

### 2.2 eMOSAIC improves OOD Binding Affinity Prediction

In the OOD setting, using structure-informed ESMFold model[43] for the protein sequence pre-training, TrustAffinity already outperforms state-of-the-art methods for the protein-ligand binding affinity predictions, as shown in Figure 2. eMOSAIC further boosts the performance of TrustAffinity by quantifying the uncertainty of predictions from it. eMOSAIC consistently outperforms the current state-of-the-art models regarding *all* four metrics in a regression task: **RMSE, MAE, Pearson Correlation**, and**Spearman Correlation**, as shown in Figure 2 A and B. eMOSAIC could successfully detect the anomalies and predict the uncertainty of predictions for improving the model performance. Although BACPI, DeepPurpose, and DeepDTA have an acceptable performance in the random split setting (Supplemental Table S1), their performances significantly drop in the scaffold split setting. In contrast, the correlation between predicted binding affinities by eMOSAIC and actual binding affinities remains high when testing chemicals have different scaffold from those in the training set. These findings clearly demonstrate the superior generalization power of eMOSAIC when predicting the binding affinity in the OOD setting. The predictions by eMOSAIC not only have higher correlation but also have significantly lower deviation as recorded by the RMSE on average 16.92%, 21.54%, and 18.97% lower, and MAE on average 20.03%, 22.55%, and 19.95% lower when compared to BACPI, DeepPurpose, and DeepDTA, respectively.

**Fig. 2.**
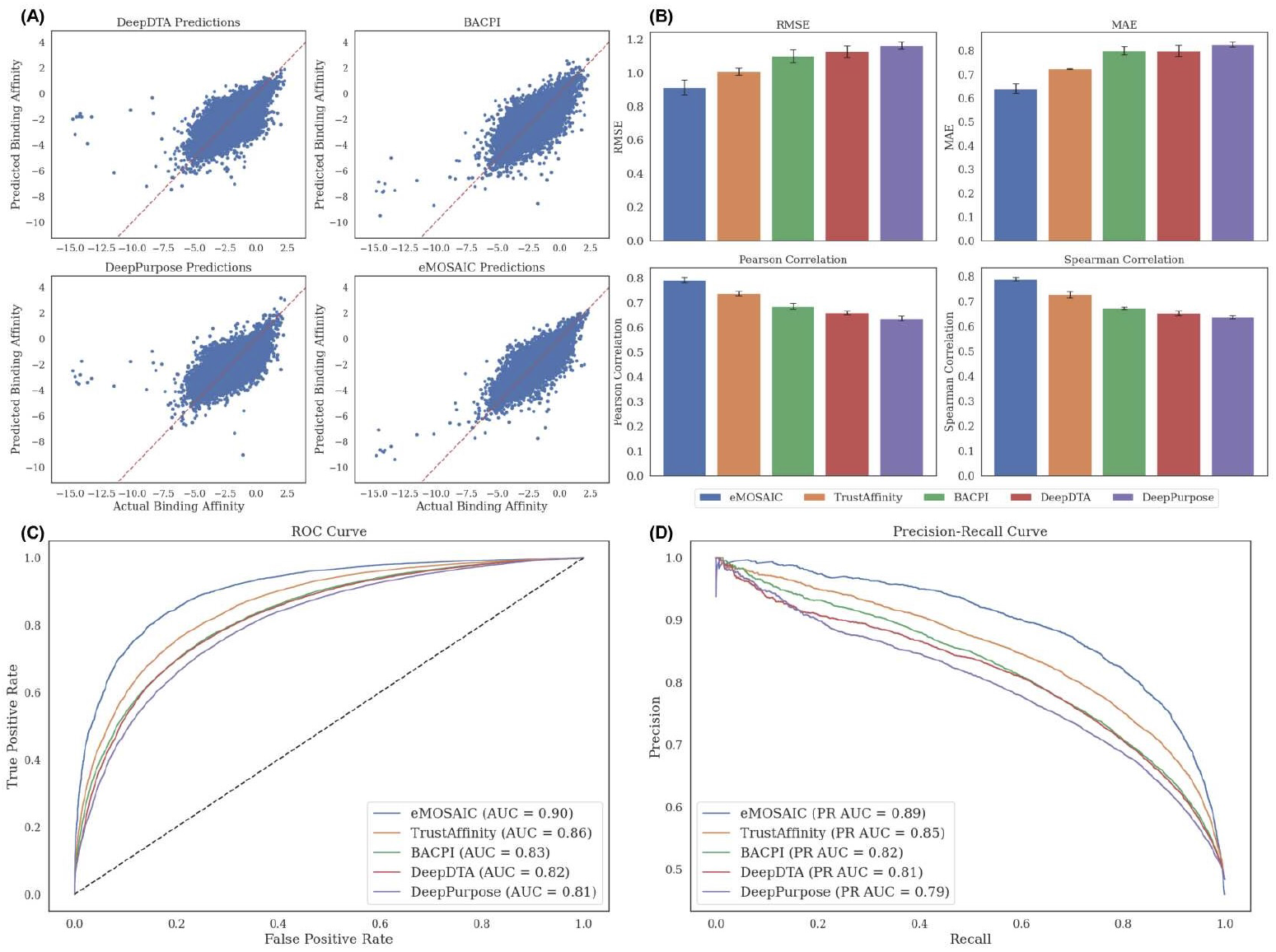
Performance comparison of eMOSAIC with baselines TrustAffinity, DeepDTA, DeepPurpose, and BACPI in the OOD setting. (A) Scatterplot of actual binding affinities vs predicted binding affinities. (B) Barplots comparing the deviations and correlations between actual binding affinities and predicted binding affinities. (C) Receiver operating characteristic (ROC) curves for the binding classification. (D) Precision-recall (PR) curves for the binding classification.

When using binding affinity greater than pKi of − 2 as a threshold, eMOSAIC also significantly outperforms all baseline models in a classification task. As shown in Figure 2 C and D, the ROC AUC and PR AUC of eMOSAIC improve 4.6% and 4.7% over TrustAffinity, respectively. The improvement over other state-of-the-art models are more significant. They are at least 9.7% and 8.5%, respectively. The performance improvement in the low false positive rate is more significant. When the false positive rate is 0.05, the positive rate of eMOSAIC is over 50% higher than the existing methods, demonstrating the potential of eMOSAIC for compound screening.

As shown in Figure 2 A, the prediction errors mainly come from the low and high affinity regions where there are few training data. Although eMOSAIC can alleviate the problem compared to other methods, further improvement is needed.

### 2.3 eMOSAIC significantly outperforms structure-based protein-ligand docking for binding affinity predictions and compound screening

We further compare the performance of eMOSAIC with PLD, which has been widely applied in compound virtual screening. When tested on the scaffold split set, PLD shows a notably poor correlation between predicted and actual binding affinities, as demonstrated in Supplemental Figure S1 and Table S2. While the structure-based deep learning model, KarmaDock, offers moderate improvement over Autodock Vina, its performance remains inadequate. In contrast, eMOSAIC significantly enhances the correlation of predicted binding affinities, showing an eight-fold improvement over KarmaDock, as shown in Figure 3. For the docking experiment, the definition of binding pockets is critical. Two methods are applied to determining binding pockets: AlphaFill approach based on the alignment with co-crystallized complex structures [51] and predicted binding pockets from a machine learning method P2Rank [52]. Although AutoDock Vina performs better with AlphaFill-predicted binding pockets compared to P2Rank’s, P2Rank outperforms AlphaFill when used with the machine learning-based method, KarmaDock.

**Fig. 3.**
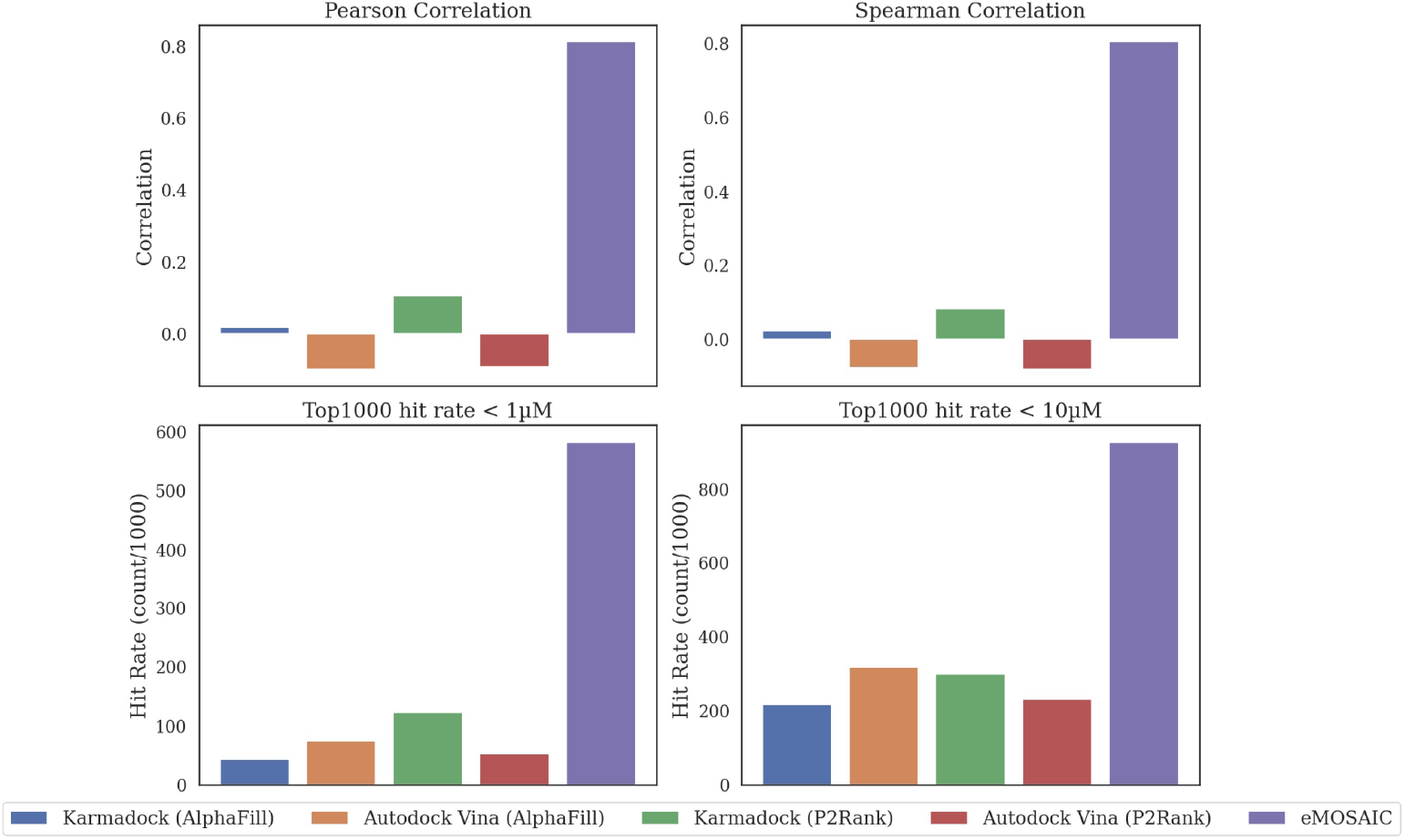
Performance comparison of eMOSAIC with structure-based methods on thePearson Correlation, Spearman Correlation, and hit rates.

For high-throughput screening applications, the ratio of high binding affinity hits in the top-*n* ranked predictions might be more interesting than the actual binding affinity. In this setting, eMOSAIC again significantly outperforms the PLD methods, as shown in Figure 3. The hit rate from eMOSAIC is approximately eight times higher than that of PLD methods for the threshold of 1 *µ*M and three and a half times higher for the threshold of 10 *µ*M. Note that a transformation is applied to align the docking score distribution with the mean and standard deviation of the ground truth pKi before computing the performance metrics. Our findings suggest that for a large-scale polypharmacology screening where understudied proteins are often involved, eMOSAIC has a clear advantage over the PLD.

Furthermore, eMOSAIC is several orders of magnitude faster than structure-based methods (Supplemental Figure S3). It takes less than 0.01 seconds for eMOSAIC, around 1 second for KarmaDock, and 30 seconds for Autodock Vina to predict the binding affinity of a protein-ligand pair. As a result, screening one million compounds against a target with eMOSAIC can be completed in just a few days.

### 2.4 eMOSAIC outperforms state-of-the-art uncertainty quantification methods

We compare eMOSAIC with three state-of-the-art methods for uncertainty quantification: RIO that is based on Gaussian process, Monte Carlo dropout, and conformal prediction. Figure 4A shows the predicted residue distribution for all predictions with actual absolute residue less than 0.5. It is clear to see that eMOSAIC has lowest upper bound of around 2.45 as opposed to the rest of the models and lowest variation. In addition, eMOSAIC has the highest Pearson Correlation between predicted residues and actual residues, as shown in Figure 4B

**Fig. 4.**
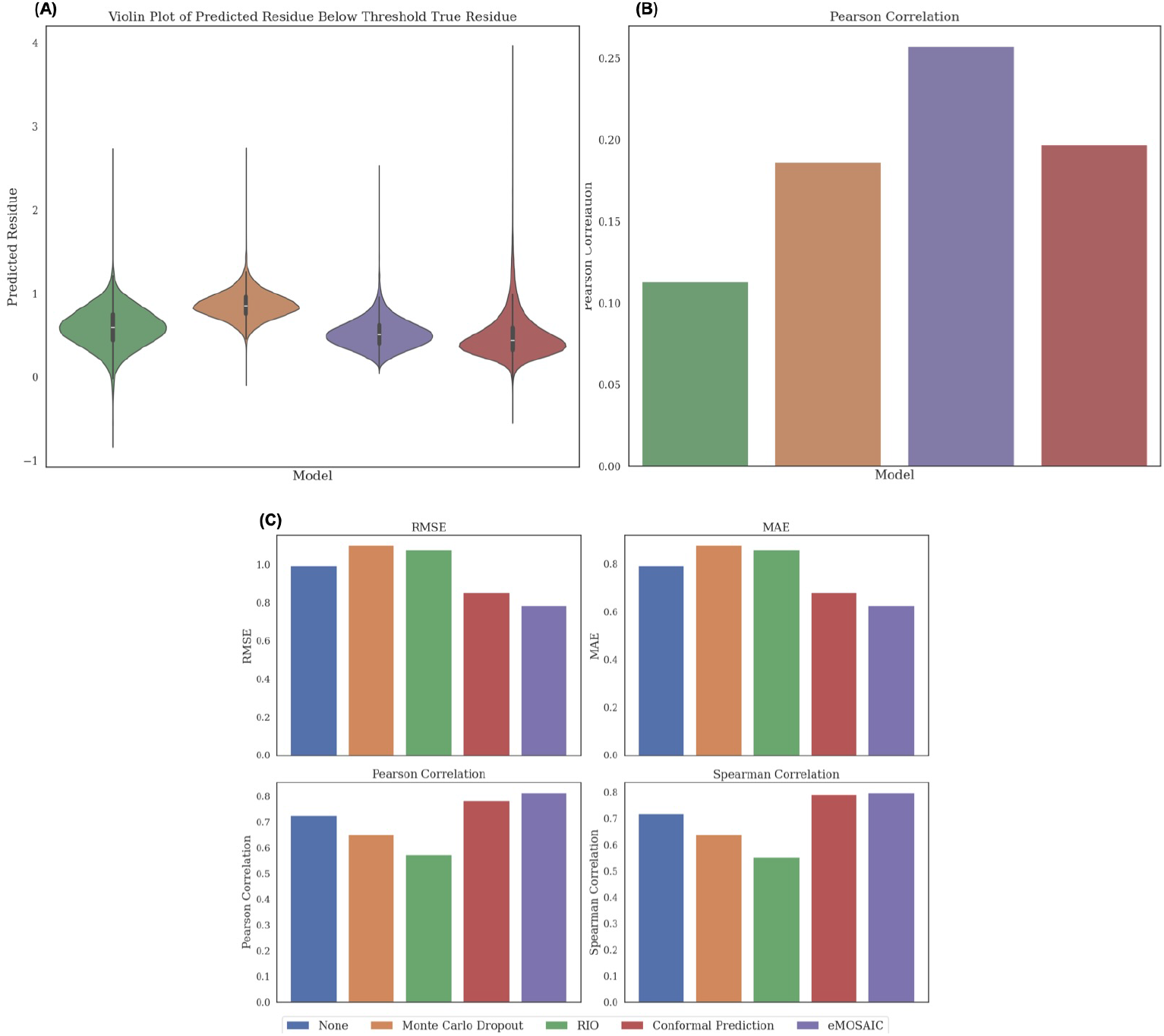
Performance comparison of eMOSAIC with state-of-the-art uncertainty quantification methods. (A) Violin Plot of predicted residue points below the high confidence threshold of 0.5. (B)Pearson Correlation of predicted residue and true residue. (C) Errors and correlations of actual and predicted binding affinities of the high confidence predictions as selected by the various uncertainty quantification models.

When evaluated for their contributions to absolute binding affinity predictions, eMOSAIC’s OOD uncertainty quantification module not only successfully identifies high-confidence, low-uncertainty PLIs but also outperforms other methods on average, as shown in Figure 4C. eMOSAIC has a 21.21% lower RMSE and 23.72% higher Pearson correlation compared to the second best method, conformal prediction. eMOSAIC can correctly identify high-confidence PLI interactions by calculating the Mahalanobis distance as an anomaly measure from the embeddings learned in the binding affinity detection module.

The GP-based RIO framework fails, due to the limited capability of the current kernels, which fail to effectively model the dependency between the PLI embeddings and residuals, resulting in less accurate residual correction and uncertainty estimation, and hence the poorest performance among all the methods.

The Monte Carlo dropout method is a crude approximation of uncertainty estimation, performing better than RIO. However, it is worse than the estimation of no uncertainty because the Bayesian posterior distribution estimated by dropout might be more complex [53]. Secondly, the Monte Carlo dropout explores limited configurations of potential weights, not all of them, resulting in incomplete uncertainty estimation [54].

Lastly, eMOSAIC performs slightly better than conformal prediction, an extensively studied method for uncertainty quantification. Interestingly, the high confident predictions from eMOSAIC and the conformal prediction is complementary, as shown in supplemental Figure S2. The Pearson Correlation between their predicted values is 0.38. It would be interesting to integrate the conformal prediction and eMOSAIC to provide a more robust statistics test for the uncertainty.

### 2.5 eMOSAIC enhances polypharmacology screening

GPCRs are compromised of targets for approximately 35% of all approved drugs, highlighting their therapeutic value. GPCRs’ druggability and accessibility make them central targets for therapeutic interventions [55]. Along with these, protein kinase inhibitors have become a critical class of drugs, especially in oncology. Up to 33% of the drug development process target these kinases [56]. Furthermore, many clinically successful therapeutics targeting GPCRs and kinases are proven to be polypharmacological drugs [10]. Therefore, we assess the models’ performance in a polypharmacological context by evaluating the ability of ligands to interact with multiple GPCRs and kinases. Predicted pKi values for PLIs are used to identify meaningful interactions. We apply a threshold of less than 100 *µ*M of the predicted pKi values, isolating strong interactions that are likely to be therapeutically significant. The evaluation is based on a multi-label classification (threshold is Ki = 100 *µ*M) to determine the potential of ligands to selectively act across different therapeutic targets and ensures that only interactions with high clinical relevance are considered. We evaluate the models using multi-label accuracy, precision, recall, F1 score, and Hamming loss. For the docking scores, a transformation was applied to align their distribution with the mean and standard deviation of the ground truth pKi before computing the performance metrics.

As shown in Table 1 the performance of eMOSAIC is significantly improved over other methods regarding all evaluation metrics for the GPCR polypharmacology. The accuracy, precision, recall, F1 score improve 21.4%, 19.7%, 20.4%, and 26.6%, respectively, while the Hamming distance reduces 164.2% compared with the second best method. For kinase polypharmacology, docking methods have the best sensitivity but poor specificity in consistent with the observations that PLD has a high false positive rate in general. eMOSAIC has a balanced sensitivity and specificity, thus it has the best accuracy and F-1 score. Among all evaluation metrics, the Hamming distance is the best for evaluating multi-label classification performance. eMOSAIC reduces the Hamming distance by 18.2% compared with the second best method.

**Table 1.** Performance comparison of polypharmacology compound screening on GPCR and Kinase Proteins. The best and the second best performed methods are in bold and underline, respectively. (\***Bold**: represents the best method, and <u>Underlined:</u> represents the second best model)

| Protein Type | Model | Accuracy ↑ | Precision ↑ | Recall ↑ | F-1 Score ↑ | Hamming Loss ↓ |
| --- | --- | --- | --- | --- | --- | --- |
| GPCR | AutoDock | 0.502 | 0.501 | 0.484 | 0.387 | 0.499 |
|  | KarmaDock | 0.51 | 0.51 | 0.143 | 0.124 | 0.49 |
|  | DeepPurpose | 0.724 | 0.696 | 0.678 | 0.668 | 0.277 |
|  | DeepDTA | <u>0.743</u> | 0.716 | 0.699 | <u>0.688</u> | <u>0.256</u> |
|  | BACPI | 0.74 | <u>0.74</u> | <u>0.722</u> | 0.652 | 0.26 |
|  | **eMOSAIC** | **0.902** | **0.886** | **0.869** | **0.871** | **0.098** |
| Kinase | AutoDock | 0.366 | 0.367 | **0.858** | 0.416 | 0.633 |
|  | KarmaDock | 0.414 | 0.414 | <u>0.735</u> | 0.38 | 0.586 |
|  | DeepPurpose | 0.698 | 0.609 | 0.637 | 0.595 | 0.302 |
|  | DeepDTA | <u>0.714</u> | 0.635 | 0.627 | <u>0.605</u> | <u>0.286</u> |
|  | BACPI | 0.681 | **0.681** | 0.407 | 0.312 | 0.319 |
|  | **eMOSAIC** | **0.758** | <u>0.662</u> | 0.648 | **0.622** | **0.242** |

## 3 Discussion

In this work, we propose eMOSAIC, a novel framework for accurate, reliable and scalable prediction of binding affinity along with an estimation of the associated uncertainty. We have successfully demonstrated the robust OOD generalization capabilities of eMOSAIC, yielding reliable (high confidence) binding affinity with high accuracy. Furthermore, we highlight the framework’s notable advantage in terms of rapid inference speed, in contrast to PLD, thereby rendering it well-suited for deployment in automated drug discovery processes to leverage uncertainty-based methodologies.

Despite eMOSAIC’s superior performance, it has certain limitations that can be further addressed. First, it lacks the ability to predict the binding pose of protein-ligand interactions (PLI), a crucial factor in the drug discovery process. eMOSAIC’s performance could be enhanced through multi-task learning, which includes predicting binding poses, binding affinity, and classifying PLI interactions. Second, the partitioning of the embedding space affects eMOSAIC’s performance. Using improved clustering methods, such as supervised contrastive learning, could yield better results. Finally, incorporating semi-supervised techniques[57] during training may boost its generalization to OOD data.

## 4 Method

### 4.1 eMOSAIC for uncertainty quantification

#### 4.1.1 eMOSAIC algorithm

Figure 1 provides an overview of eMOSAIC. We utilize embedding clustering and Mahalanobis distance to identify anomalies and quantify uncertainties. P.C. Mahalanobis introduced the metric known as Mahalanobis Distance (MD) in 1936 as a measure of anomaly and for detecting outliers [58]. For a given point in a distribution, the MD is defined as follows:

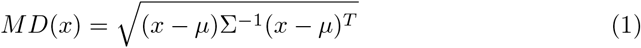

Here, *µ* and Σ refer to the mean and variance of the distribution. It considers the variance between the various variables in a multivariate distribution since real-life data often contains many correlated variables. Mahalanobis distance has normalization through division by the covariance matrix, ensuring that variables with different scales are suitably handled. Because of these properties, it can effectively identify outliers or anomalies from the main distribution. Mahalanobis distance has previously been used in deep learning frameworks for anomaly, OOD, and adversarial detection in computer vision, time series, and natural language processing tasks [59–62]. These instances have clearly motivated the usage of Mahalanobis distance for anomaly detection tasks to improve the reliability of predictions by deep learning frameworks.

After training of a deep learning model, a binding affinity module in this paper, we extract these attentive pooling embeddings which represent the PLI. Now we perform k-means clustering of the PLI embeddings from the training set, and for each cluster we obtain the mean (*µ*) and variance matrix (**Σ**) for Mahalanobis distance calculation. Given an embedding of a unseen PLI, we calculate the Mahalanobis distance to each cluster. Then we use these distances obtained as features to train a simple multi-layer perceptron (MLP) to predict the absolute residue, which is defined as the difference between the predicted binding affinity and the true binding affinity. This way the MLP learns to identify the PLI anomalies as predicted by the model. For outlier detection, we select points whose predicted residue is less than 0.5, ensuring that only high-confidence predictions are filtered.

#### 4.1.2 Training of eMOSAIC

We first train the binding affinity prediction module. After training, we obtain the embeddings with the best model based on the validation set, and train the OOD uncertainty quantification module to identify outliers. More details about the hyperparameters and configuration of the eMOSAIC Model are present in the appendix.

### 4.2 Binding Affinity Module

The binding affinity module consists of three sub-modules - the protein sequence module, the ligand processing module, and lastly the PLI module. All of these modules work together to predict the binding affinity associated with the PLI.

#### 4.2.1 Protein Sequence Module

Protein sequence representation is one of the most vital components in the machine learning frameworks for predicting not only PLIs [34, 45, 46, 63], but also their 3D structure [23, 43]. Protein sequences contain information that can be used to infer protein structure, function, and family [43], making them a rich source of data for machine learning models [23, 43]. Large datasets of protein sequences are available [23, 64, 65], enabling machine learning frameworks to learn high-level, general representations of proteins. We utilize ESMFold [43] to obtain the protein sequence embeddings, which deploys a large language model (LLM) - ESM-2 alongside a folding module and a structure module for modeling the protein structure. The ESM-2 protein language model, which is able to capture the protein structures at the fine resolution of the atomic level, consists of variable parameters ranging from 8M to 15B. We use the 650M parameter model to obtain the refined protein sequence representation. We observed that the sequence representations obtained from the structure module of the ESMFold model performed better than the protein embeddings obtained from the ESM-2 model directly as well as the sequence embeddings obtained from the folding block, possibly because the structure block refines the protein sequence obtained from the ESM-2 model. We remove protein sequences greater than 700 in length as they are very low in numbers and due to constraints with time and memory. Since the embeddings obtained are variable in size corresponding to the protein sequence length, to make them consistent for the next steps, we perform padding to pad the sequences with lengths less than 700, and define masks associated with the sequences that track the padding. Since CNNs are known to work well with processing sequence representations, we use the ResNet model [66] with 5 layers, and each layer has 4 convolutional layers to obtain a refined protein sequence embedding. Finally, adaptive masking (using interpolation) is used based on the changes to the embeddings to avoid the loss of information.

#### 4.2.2 Ligand Module

We represent each ligand as a 2D graph, where the nodes symbolize atoms and the edges are bonds. Embeddings for both node and edge are learnt using the graph isomorphism network (GIN) [67]. For atom or node attributes, we used atom types, hybridization types, atom degrees, atom chirality, atom formal charges and atom aromatic all converted to one-hot encoding before being utilized by GIN. We use a 5-layer GIN architecture, which aggregates and updates node embedding for each atom/node. To obtain a graph-level or a ligand-level embedding that remains permutation invariant, a final sum pooling operation is used.

#### 4.2.3 PLI Module

After obtaining both the protein and ligand embeddings, we use the attentive pooling network such that the model is aware of both protein and ligand and that the interaction isn’t solely dependent on either of protein or ligand. This network gives us the attention weighted embeddings for both which are then concatenated and fed to a MLP which predicts the final binding affinity.

### 4.3 Experiments

#### 4.3.1 Dataset

We train TrustAffinity on the ChEMBL31 database [30], which consists of 350400 PLI pairs. In the experiments, we split the dataset into training, testing, and validation set by 7:2:1. Negative log transformation was performed on Ki (binding affinity) to obtain pKi values. The data was split using the following methods - 1) Random Split - random selection of protein-ligand pairs, 2) Random Scaffold Split - random selection of scaffolds of chemical structures[68] such that the chemicals in the testing set have different scaffolds from those in the training/validation set. Scaffold split ensures that there was no overlap of scaffold in the training, testing and validation set. This was done to validate the model’s generalization power in real-world OOD setting. Moreover, considering the vast chemical space for drug discovery, it is very likely that the model will encounter unknown and new scaffolds.

#### 4.3.2 Baseline models

We test eMOSAIC against baselines in two different objectives, 1) Binding Affinity Prediction and 2) Uncertainty Quantification.

For binding affinity prediction task we compare eMOSAIC with both sequence-based deep learning models such as BACPI [45], DeepDTA [46], DeepPurpose [47] as well as a typical PLD method, AutoDock Vina [48], and deep learning-based docking model KarmaDock [49] on the OOD test set. In order to compare with docking programs, AutoDock and KarmaDock were used to predict the docking scores for ligand-protein pairs for the scaffold splitting data set. Alphafill [51] annotated binding pockets were used to define the searching space for AutoDock and KarmaDock. After removing the binding pockets for 38 different irons, such as FE, NA, 1246 binding pockets on 429 proteins were used as the pre-defined binding pockets to set up docking. If there are multiple pockets on one protein, the ligand will be docked into all pockets and the one with the best docking score will be selected for this ligand-protein pair. For other proteins not in Alphafill dataset, P2Rank [52] was used to predict the binding pockets on their Alphafold predicted model structures. The ligands were then docked into these binding pockets by AutoDock and KarmaDock and the best docking scores were selected for these ligand-protein pairs. In order to evaluate how many of the top ranked pairs have high binding affinities, top 1000 hit rates were calculated, which measures how many of the top 1000 ranked pairs have pKi *>* -log(1) or -log(10). A higher top_1000_ hit rate correlates with better docking rank performances.

Along with this, we also compare the performance of our uncertainty quantification module with other uncertainty quantification methods including GP-based RIO framework [36], Monte Carlo Dropout [50], and Conformal Prediction [38].

Conformal prediction relies on obtaining a nonconformity score to measure the confidence interval for the predictions. This nonconformity score is crucial, and several different methods exist to compute it [39], and several methods are available that describe various variants of Conformal Prediction [69]. The most common nonconformity measures for regression-based models are based on absolute error, including using the calibration set’s absolute errors to provide intervals for the new predictions and using predicted absolute errors [39]. In our case, we use the attentive pooling embeddings to train an MLP for predicting the absolute error in case of CP baseline. We then select the points which have a lower than average length of confidence interval in the test set. We do the same for obtaining the high confidence points in case of Monte-Carlo Dropout method.

#### 4.3.3 Evaluation

We evaluate eMOSAIC’s performance using root mean square (RMSE), mean absolute error (MAE), Pearson correlation coefficient (*r*) and Spearman’s rank correlation coefficient (*ρ*).

## Notes

### Competing Interest Statement

The authors have declared no competing interest.

### Summary of Updates

Extensive analysis and more baselines comparison.

